# Producing Hfq/Sm Proteins and sRNAs for Structural and Biophysical Studies of Ribonucleoprotein Assembly

**DOI:** 10.1101/150672

**Authors:** Kimberly A. Stanek, Cameron Mura

**Author notes:** Correspondence can be addressed to KS or CM.

## Abstract

Hfq is a bacterial RNA-binding protein that plays key roles in the post–transcriptional regulation of gene expression. Like other Sm proteins, Hfq assembles into toroidal discs that bind RNAs with varying affinities and degrees of sequence specificity. By simultaneously binding to a regulatory small RNA (sRNA) and an mRNA target, Hfq hexamers facilitate productive RNA⋯RNA interactions; the generic nature of this chaperone-like functionality makes Hfq a hub in many sRNA-based regulatory networks. That Hfq is crucial in diverse cellular pathways—including stress response, quorum sensing and biofilm formation— has motivated genetic and ‘RNAomic’ studies of its function and physiology (*in vivo*), as well as biochemical and structural analyses of Hfq⋯RNA interactions (*in vitro*). Indeed, crystallographic and biophysical studies first established Hfq as a member of the phylogenetically-conserved Sm superfamily. Crystallography and other biophysical methodologies enable the RNA-binding properties of Hfq to be elucidated in atomic detail, but such approaches have stringent sample requirements, *viz*.: reconstituting and characterizing an Hfq•RNA complex requires ample quantities of well-behaved (sufficient purity, homogeneity) specimens of Hfq and RNA (sRNA, mRNA fragments, short oligoribonucleotides, or even single nucleotides). The production of such materials is covered in this Chapter, with a particular focus on recombinant Hfq proteins for crystallization experiments.

**Abbreviations:**

3D: three-dimensional
AU: asymmetric unit
CV: column volume
DEPC: diethyl pyrocarbonate
HDV: hepatitis δ virus
HDVD: hanging-drop vapor diffusion
IMAC: immobilized metal affinity chromatography
MW: molecular weight
MWCO: molecular weight cut-off
nt: nucleotide
PDB: Protein Data Bank
RNP: ribonucleoprotein
RT: room temperature
SDVD: sitting-drop vapor diffusion

**Journal format:** *Methods in Molecular Biology* (*Springer Protocols* series); this volume is entitled “*Bacterial Regulatory RNA: Methods and Protocols*”; an author guide is linked at http://www.springer.com/series/7651

## 1 Introduction

The bacterial protein Hfq, initially identified as a host factor required for the replication of bacteriophage Q³ RNA [1], plays a central role in RNA biology: both in RNA-based regulation of gene expression and in modulating RNA stability and lifetime *in vivo* [2]. Hfq functions broadly as a chaperone, facilitating contacts between small non-coding RNAs (sRNAs) and their cognate mRNAs [3]. The RNA interactions may either stimulate or inhibit expression, depending on the identity of the mRNA–sRNA pair and the molecular nature of the interaction (high or low affinity, stable or transient, etc.) [4,5]. In many cases, Hfq is required for these pairings to be effective [6], and knockdown of the *hfq* gene results in pleiotropic phenotypes such as increased UV sensitivity, greater susceptibility to oxidative or osmotic stress, decreased growth rates, etc. [7]. A flood of ‘RNAomics’-type studies, over the past decade, has shaped what we know about Hfq-associated RNAs [2,8,9]. Hfq has been linked to many cellular pathways that rely on rapid responses at the level of post-transcriptional/mRNA regulation, including stress responses [10-12], quorum sensing [13], biofilm formation [14], and virulence factor expression [15,12].

Hfq homologs are typically ≈80-100 amino acids in length, with the residues folding as an α-helix followed by five β-strands arranged into a highly-bent, antiparallel β-sheet [16,17]. Hfq monomers self-assemble into a toroidal hexamer, the surface of which features at least three distinct regions that can bind RNA. The *proximal* face of the hexamer (proximal with respect to the *N*′-terminal α-helix) is known to bind U-rich sequences [16,18], while the *distal* face of the (Hfq)_6_ ring binds preferentially to A-rich RNA elements [19,20]. Recently a third, lower-affinity, *lateral* surface on the outer rim of the Hfq ring has been shown to bind RNA [21] and aid in sRNA…mRNA annealing [22]. This lateral site likely has a preference for U-rich segments [23], but also may interact fairly non-specifically with RNA because of an arginine-rich region that is found in some homologs. While the exact mechanism by which Hfq facilitates productive RNA…RNA interactions remains unclear, it is thought that the distal face binds the 3′-poly(A) tails of mRNAs while the proximal face binds to 3′ U-rich regions of sRNAs [3]. The lateral surface may act either to cycle different RNAs onto Hfq [22] or as an additional surface for binding to internal, U-rich regions of an sRNA. A recent study suggested that this mechanistic model holds for only a subset of sRNAs, termed ‘Class I’ sRNAs [24]. A second subset of sRNAs (‘Class II’) appears to bind both the proximal and distal sites of Hfq; the mRNA targets of these Class II sRNAs are predicted to bind preferentially to the lateral region of the Hfq ring.

A detailed understanding of how different sRNAs interact with Hfq, and with target RNAs in a ternary RNA•Hfq•RNA complex, requires atomic-resolution structural data. While multiple structures have been determined for short (≲10-nucleotide) RNAs bound to either the proximal [16,18], distal [19,20], or lateral [23] sites of Hfq (Table 1), as of this writing only one structure of an Hfq bound to a full-length sRNA has been reported [25]. In that Hfq•RNA complex, comprised of *E. coli* Hfq bound to the *Salmonella* RydC sRNA (Fig 1A), the 3′-end of the sRNA encircles the pore, towards the proximal face of the hexamer, while an internal U–U dinucleotide binds in one of the six lateral pockets on the periphery of the Hfq ring (Fig 1B). Though other regions of the sRNA were found to further contact a neighboring Hfq ring in the lattice (Fig 1C), the stoichiometry of the Hfq•RydC complex *in vivo*, at limiting RNA concentrations, is thought to be 1:1. (Interestingly, two distinct interaction/binding modes were seen between RydC and the lateral rim of an adjacent hexamer [Fig 1C].) The Hfq•RydC complex offers a valuable window into our understanding of Hfq…sRNA interactions, limited mainly by the relatively low resolution (3.48 Å) of the refined structure. For this and other Hfq•RNA complexes, many questions can be addressed by leveraging different types of structural and biophysical approaches. Ideally, the methods used would provide a variety of complementary types of information (i.e., the underlying strategy in taking a ‘hybrid methods’ approach [26,27]).

**Table 1:**
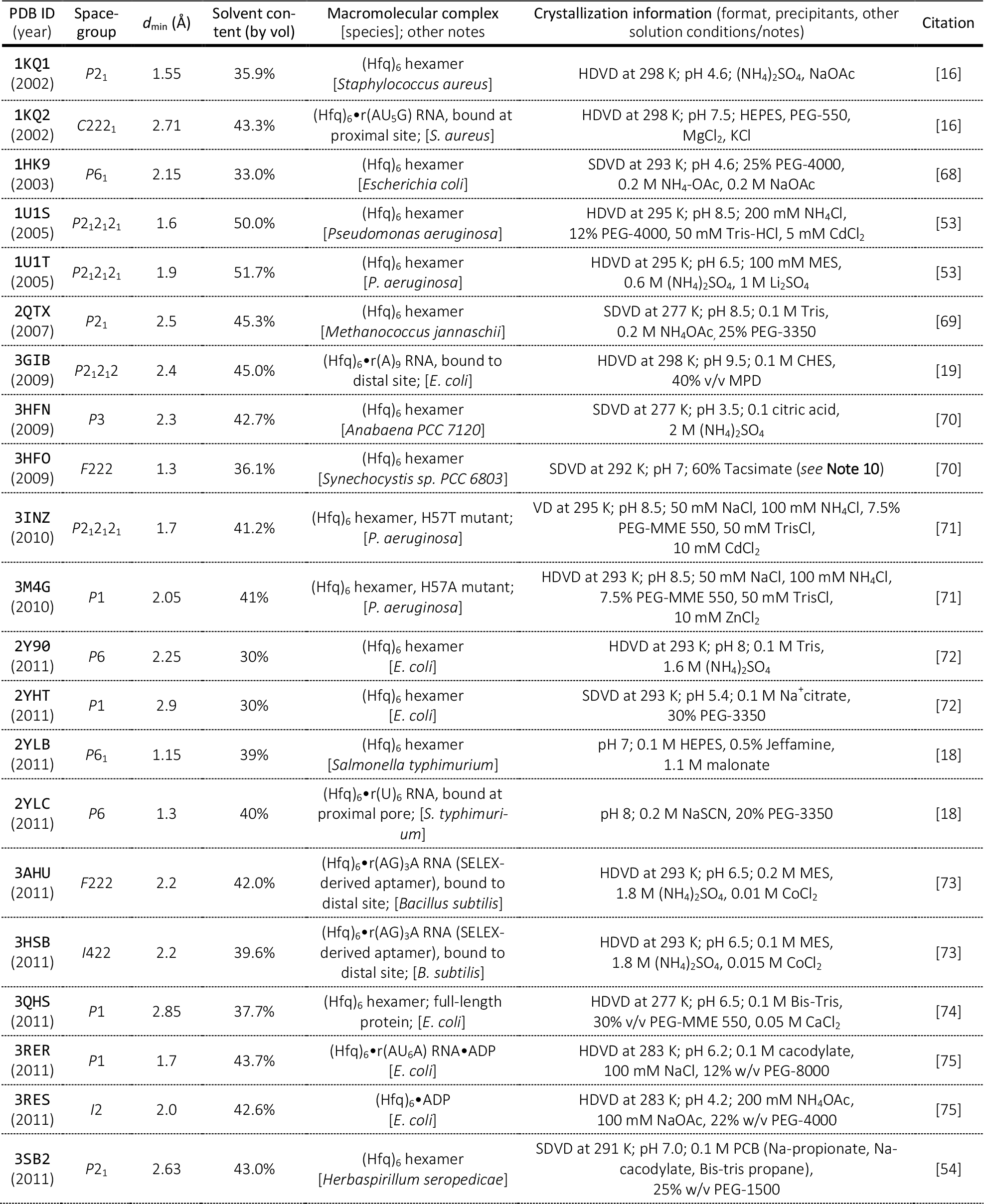
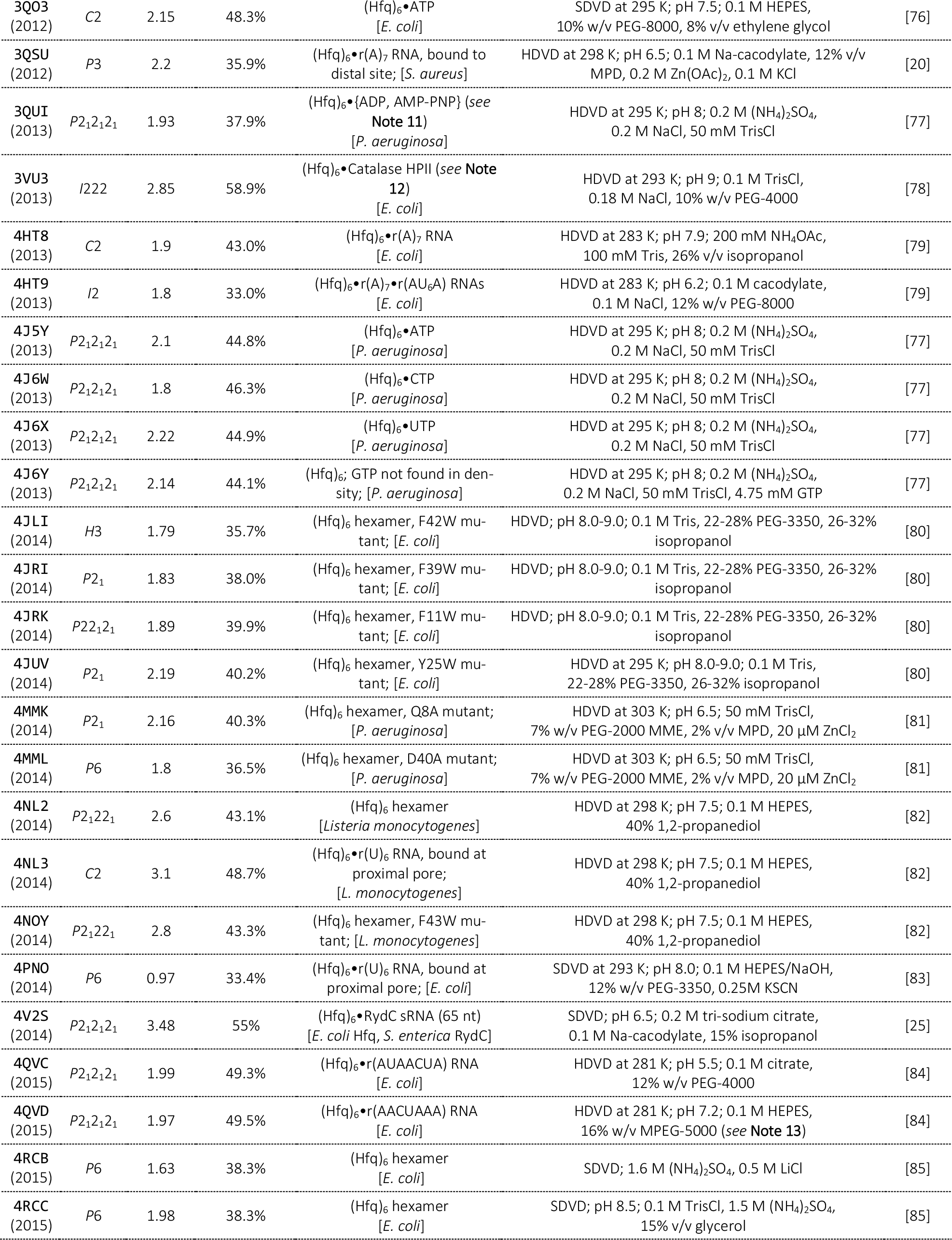
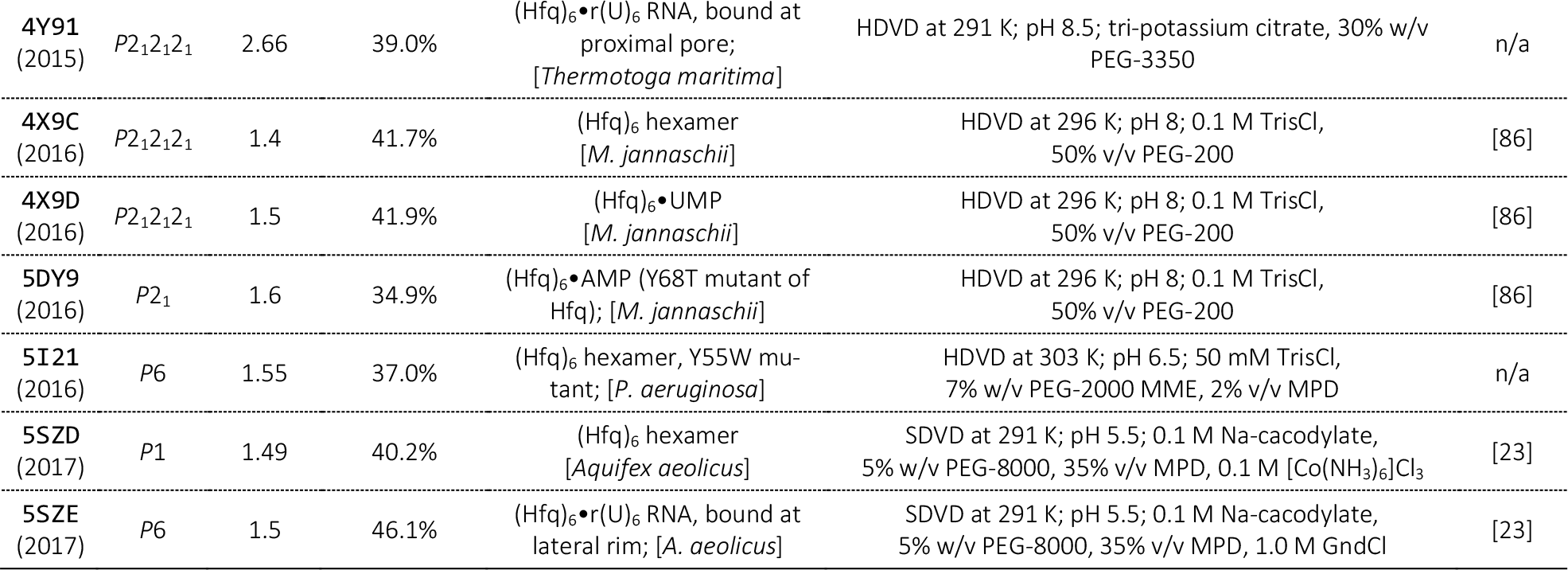
A comprehensive list of Hfq structures in the PDB, including co-crystal structures with nucleotides and RNAs.

Atomic-resolution information may be obtained in several ways. Historically, the premier methodologies have been X-ray crystallography and solution NMR spectroscopy; these well-established approaches are described in many texts, such as [28] and [29]. Though beyond the scope of this chapter, note that much progress in recent years has positioned electron cryo-microscopy (cryo-EM) as a powerful methodology for high-resolution (nearly atomic) structural studies of macromolecular assemblies[30,31], including ribonucleoprotein (RNP) complexes such as the ribosome [32-34], telomerase [35] and, most recently, the spliceosome [36,37]. Thus far, all Hfq and Hfq•RNA structures deposited in the Protein Data Bank (PDB), listed in Table 1, have been determined via X-ray crystallography. The molecular weight (MW) of a typical Hfq hexamer is ≈60 kDa while sRNAs, which range in length from ≈50–500 nucleotides (nt), have MWs of ≈16–1600 kDa. An RNP complex of this size is ideally suited to macromolecular crystallography.

In this Chapter, we describe how to prepare and crystallize Hfq•sRNA complexes for structure determination and analysis via the classic, single-crystal X-ray diffraction approach. However, if crystallographic efforts with a particular Hfq or Hfq•sRNA complex prove difficult because of a lack of well-diffracting crystals—or, even when such crystals can be reproducibly obtained—then one can also consider investigating the Hfq-based complex via complementary approaches. Two main families of alternative methodological approaches are available: (i) NMR and other spectroscopic methods (e.g., EPR [38]), and (ii) cryo-EM and other scattering-based approaches (e.g., SAXS [39]). NMR and cryo-EM are routinely used for smaller or larger-sized biomolecular complexes, respectively, though methodological developments are continuously redefining these limitations. The current upper size limit for *de novo* NMR structure determination is ≈40 kDa, this limit being reached via the application of techniques such as TROSY, as well as relatively recently developed approaches for deriving distance restraints (e.g., paramagnetic relaxation enhancement). NMR applications to RNP complexes have been recently reviewed [40]. In the reverse direction, from large to small, cryo-EM was recently used to determine the structure of a protein as small as ≈170 kDa [41]; the highest-resolution cryo-EM structure reported thus far has reached a near-atomic 2.2 Å [42].

As alluded to above, RNA and RNP complexes pose particular challenges in crystallographic structure determination [43,44]. Most proteins typically adopt a discrete, well-defined three-dimensional (3D) structure, but populations of RNAs tend to sample broad ranges of conformational states, yielding greater structural heterogeneity; notably, this holds even if the sample is technically monodisperse (i.e., homogeneous in terms of MW). Therefore, RNA and RNP crystals often exhibit significant disorder and correspondingly poorer diffraction, as gauged by resolution, mosaicity, and other quality indicators [45]. Synthesis and purification of an RNA construct through *in vitro* transcription (see §3.2) generates large quantities of chemically homogeneous RNA, and also conveniently lends itself to the engineering of constructs that may be more crystallizable, or that exhibit improved diffraction. Alongside crystallization efforts, chemical probing [46] and structure prediction/modelling methods [47] can be used to examine the secondary structure of the RNA of interest, as well as identify potential protein-binding sites. Then, in designing a more crystallizable construct, extraneous regions of RNA can be either removed or replaced with more stable secondary structures (e.g. stem-loops incorporating tetraloop/tetraloop-receptor pairs); these rigid structural elements can aid crystal contacts and enhance lattice order [44,48]. As an example of judiciously choosing (and/or designing) an RNA system for crystallographic work, the aforementioned RydC sRNA (Fig 1A) is a favorable candidate for crystallization efforts because (i) it is relatively small and compact (forming a pseudoknot), and (ii) it features multiple U-rich regions that can potentially bind to both its cognate Hfq (within a single RNP complex) as well as other Hfq proteins across the lattice. In the crystal structure (Fig 1B, C), the RNA was found to span two Hfq hexamers, forming intermolecular contacts that helped stitch together a stable crystal lattice.

Once crystals that diffract to even low resolution (e.g., ≲ 4 Å) are obtained, a native (underivatized) X-ray diffraction dataset can be collected. From this dataset alone, much can be learned [49], including the likely stoichiometry in the specimen that crystallized (e.g., 1:1 or 2:1 Hfq:RNA?) and whether or not the complex found in the crystalline asymmetric unit (AU) features any additional (non-crystallographic) symmetry. Calculation of an initial electron density map from the diffraction data requires approximate phases for each X-ray reflection. Such phases can be estimated, *de novo*, via a family of computational approaches based on the two fundamental ideas of multiple isomorphous replacement (MIR) or multi-wavelength anomalous dispersion (MAD); these general approaches require derivatization of native crystals, either via soaking with heavy atoms (MIR/SIR/etc.) or covalent introduction of an anomalously scattering atom (MAD/SAD/etc.) such as selenium. For more information on approaches to *de novo* estimation of initial sets of phases, see [28].

If a known 3D structure is similar to the (unknown) structure that one seeks to determine, then the phase problem can be greatly simplified. In such cases, a phasing approach known as molecular replacement (MR) can be used to estimate initial phases for the unknown structure (the ‘target’) using a known structure (the ‘probe’), or a suitable modification thereof (e.g., a homolog model). Essentially, the MR approach can be thought of as a ‘fit’, via rigid-body transformations that sample the three rotational and three translational degrees of freedom, of the probe structure to the unknown phases of the diffraction data (which, in turn, directly result from the detailed 3D coordinates of the target structure). Because 3D structures are available for Hfq homologs from many species (Table 1), the phase problem is much simplified by using MR. Similarly, the phases computed in refining a given *apo* Hfq structure can then be used as an initial phase estimate for X-ray data collected for a corresponding Hfq•sRNA complex. The protocols below assume that one can successfully estimate initial phases via MR; if such is not the case, e.g., if there is an unexpected/complicated stoichiometry in the AU (say four Hfq rings and three RNAs, in an odd geometric arrangement), then one must resort to *de novo* phasing methods.

## 2 Materials

### 2.1 Hfq purification

1. DNA sample that contains the *hfq* gene of interest (e.g., genomic DNA)
2. Inducible expression vector capable of encoding a His6x affinity tag (e.g., pET-22b(+) or pET28b(+))
3. Chemically competent BL21(DE3) *E. coli* cells
4. Lysogeny broth (LB) agar plate, supplemented with appropriate antibiotic (e.g., 100 ¼g/ml ampicillin or 50 ¼g/ml kanamycin)
5. LB liquid media, supplemented with a suitable antibiotic (e.g., 100 ¼g/ml ampicillin or 50 ¼g/ml kanamycin)
6. 1 mM isopropyl *β*-D-1-thiogalactopyranoside (a 1000× stock [1 M] can be prepared, partitioned into 1-ml aliquots, and stored at –20 °C)
7. Lysis buffer: 50 mM Tris pH 7.5, 750 mM NaCl
8. Chicken egg-white lysozyme (100x stock at 1 mg/ml)
9. 0.2-¼m syringe filters
10. His-Trap HP pre-packed sepharose column (GE Healthcare)
11. 200 mM Ni_2_SO_4_
12. Wash buffer: 50 mM Tris pH 7.5, 200 mM NaCl, 10 mM imidazole
13. Elution buffer: 50 mM Tris pH 7.5, 200 mM NaCl, 600 mM imidazole
14. Bovine thrombin (200 U/ml)
15. p-aminobenzamidine-agarose resin (Sigma)
16. 3-kDa molecular weight cut-off (MWCO) filter-concentrators (Amicon)

### 2.2 sRNA purification

1. Template DNA (≈1 μg/μl for a 3-kb linearized plasmid)
2. 10x transcription buffer: 500 mM Tris-HCl pH 8.0, 100 mM NaCl, 60 mM MgCl_2_, 20 mM spermidine
3. 10× rNTP mix: rATP, rCTP, rUTP, rGTP, each at 20 mM
4. 100 mM dithiothreitol (DTT)
5. Diethyl pyrocarbonate (DEPC)–treated H_2_O (RNase-free)
6. T7 RNA polymerase (20 U/μl)
7. RNase-free DNase I (50 U/μl)
8. Denaturing (8 M urea) 5% polyacrylamide gel
9. Elution buffer: 20 mM Tris-HCl pH 7.5, 1 mM EDTA, 1% (w/v) SDS
10. Phenol:chloroform (1:1)
11. 96% ethanol

### 2.3 Co-crystallization trials

1. Purified Hfq protein (see §3.1)
2. Purified sRNA construct (see §3.2)
3. The ‘Natrix’ and ‘Crystal Screen’ sparse-matrix crystallization screens (Hamton Research)
4. Intelliplate 96-3 Microplates (Hampton Research)
5. Sealing tape (Hampton Research)
6. 24-well VDX plates with sealant (Hampton Research)
7. Siliconized glass cover slips (Hampton Research)
8. Vacuum grease
9. Light stereomicroscope, with cross-polarizing lenses (e.g., Zeiss Discovery V20)

## 3 Methods

### 3.1 Hfq Purification

Crystallization efforts typically require large quantities of highly-purified and concentrated material, e.g., on the scale of >100 μl at >10 mg/ml of the biomolecule. To achieve this, Hfq is often expressed in a standard *E. coli* K12 laboratory strain, using a plasmid-based construct created via standard recombinant DNA techniques. The expression vector, e.g. an inducible T7*lac*-based system (pET series), can be used to add various affinity tags to the N′ or C′ termini of the wild-type sequence. Then, over-expressed Hfq can be readily purified via affinity chromatographic means. Further steps, detailed below and in Fig 2, may be required to remove co-purifying proteins and nucleic acids. Because at least some Hfq homologs bind nucleic acids fairly indiscriminately, the ratio of absorbances at 260 nm and 280 nm (A_260_/A_280_) should be monitored over the various stages of purification in order to detect the presence of contaminating nucleic acids. Pure nucleic acid is characterized by an A_260_/A_280_ ratio of ≈1.5-2.0, versus ≈0.7 for pure protein; rapid colorimetric assays can be used alongside absorbance readings to discern whether a contaminant is mostly RNA or DNA [50].

In terms of solution behavior, experience has shown that Hfq homologs generally remain soluble in aqueous buffers at temperatures >70 °C, and that they resist chemical denaturation (e.g., treatment with 6 M GndCl); in many cases, depending on the species of origin, Hfq samples are insoluble at low temperatures. We have found that the common practice of purifying/storing proteins at 4 °C can be unwise with Hfq homologs: if visible precipitation occurs at ≈4 °C, we recommend that purification be conducted at ambient room temperature (≈18–22 °C), and that elevated temperatures be considered for long-term storage of purified protein (e.g., ≈37–42 °C works well for an Hfq homolog from the hyperthermophile *Aquifex aeolicus* ). In terms of protein expression behavior, purification strategies, solubility properties (temperatureand ionic strength–dependence), etc., we have found that the *in vitro* behavior of many Hfq constructs resembles the overall properties of Hfq orthologs (Sm proteins) from the archaeal domain of life [17].

Previously, His-tagged [51,18] and self-cleaving intein-tags [52,16,21] have been used for affinity purification of Hfq, although it has also been purified without the use of a tag. Un-tagged Hfq has been purified using immobilized-metal affinity chromatography (IMAC), as the native protein has been shown to associate with the resin [4]. Poly(A)-sepharose [10] and butyl-sepharose [53,54] columns also have been utilized to purify untagged Hfq, leveraging the RNA-binding properties and partially hydrophobic nature of the surface of the protein (respectively). Below, we outline the purification of recombinant Hfq using a His6x-tagged construct. This tag can be removed at a later step through protease treatment; this is a crucial feature, as it is possible that even a modestly-sized His6x tag can interfere with the oligomerization behavior and binding properties of Hfq [55]. The <u>i</u>ntein-<u>m</u>ediated <u>p</u>urification with an <u>a</u>ffinity <u>c</u>hitin-binding <u>t</u>ag (IMPACT) scheme, used to both clone and purify intein-tagged constructs, has been successfully applied to Hfq by multiple labs (e.g., [52,21]). This system is available as a kit from NEB, so that method will not be described herein. Note that many of the considerations and notes described below, for His6×-tagged Hfq, also apply when purifying and working with any Hfq construct, intein–based or otherwise.

1. Clone the *hfq* gene, using a genomic sample as PCR template, into an appropriate expression vector; ideally, such a vector will add a His-tag. A compatible vector from the pET series of plasmids often works well (e.g., pET-28b(+), which fuses an *N*-terminal His6× tag).
2. Transform competent BL21(DE3) *E. coli* with the recombinant Hfq plasmid and plate onto LB agar supplemented with antibiotic (e.g., 50 μg/ml kanamycin if using pET-28b(+)).
3. Grow-out the transformed cells in LB media at 37 °C with shaking (225 rpm) to an optical density at 600 nm (OD_600_) of ≈0.6-0.8. Then, induce over-expression of Hfq by adding IPTG to a final concentration of 1 mM. Optionally, immediately before adding IPTG take a 1-ml aliquot of the cell culture as a *t* = 0 (pre-induction) sample; this sample can be stored at –20 °C and later analyzed alongside a post-induction sample (by SDS-PAGE) in order to assess overexpression levels.
4. Incubate the cell cultures for an additional 3-4 h at 37 °C, with continued shaking, and then centrifuge at 15000*g* for 5 min to pellet. Optionally, take a 1-ml aliquot of the cell culture at *t* ≈2–3 h post-induction; this sample can be stored at –20 °C and analyzed by SDS-PAGE.
5. Resuspend the cell pellet from the previous step in Lysis Buffer (§2.1). Optionally, DNase I and RNase A can be added at this stage in order to hydrolyze any nucleic acids, Hfq-associated or otherwise (*see* **Note 1**).
6. Incubate the lysate with 0.01 mg/ml lysozyme for 30 min (if a more thorough chemical lysis is required), at either RT or 37 °C; gently shake/invert the sample a few times during this incubation.
7. Mechanically lyse the cells using a sonicator or other similar means (e.g., a microfluidizer or French press). Remove cellular debris from the lysate by centrifugation at 16000*g* for 5-10 min at RT.
8. Initial purification of Hfq can be achieved by using a heat-cut to precipitate endogenous, mesophilic *E. coli* proteins. Proceed by incubating the supernatant from the last step (i.e., clarified lysate) at ≈70–80 °C for ≈10–15 min (*see* **Note 2**); a substantial amount of white precipitate should develop within minutes. Next, use a high-speed centrifugation step (e.g., 33000*g* for 30 min) to clarify the soluble, Hfq-containing supernatant; for pilot studies, the supernatant and pellet fractions from this step can be saved in case SDS-PAGE analysis becomes necessary.
9. If nucleic acid is still present in the heat-treated sample, as assessed by A_260_/A_280_ ratios, colorimetric assays [50], or dye-binding assays (e.g., cyanine-based stains such as PicoGreen or SYBR-Gold), then a chaotropic agent such as urea or GndCl can be added to the sample, to a concentration of up to 8 M or 6 M, respectively (*see* **Note 3**).
10. Pass the latest Hfq-containing sample through a 0.2-μm filter (syringe or vacuum line) to remove any particulate matter, prior to applying the material to a high-performance liquid chromatography (HPLC) or fast protein liquid chromatography (FPLC) system in the next step.
11. Isolate the His6×-tagged Hfq via IMAC, using an iminodiacetic acid sepharose resin in a prepacked column connected to an HPLC or FPLC instrument. All buffers should be vacuum-filtered (0.45-μm filters) and sonicated before use. In brief, this IMAC step entails the following sub-steps:

1. Prepare the resin by washing with 3-4 column volumes (CVs) of dH_2_O, and then 3-4 CVs of Wash Buffer.
2. Charge the resin with Ni^2^+ by loading at least 1-2 CVs of 200 mM NiSO_4_ (*see* **Note 4**).
3. Load the crude (unpurified) Hfq-containing protein sample (collect the flow-through), and then wash the column with several CVs of Wash Buffer (until the A_280_ trace drops near baseline).
4. Elute the Hfq protein by applying a linear gradient of Elution Buffer, from of 0→100% over 10 CVs.
5. Combine the fractions thought to contain Hfq (as assessed by A_280_ and the elution profile), and dialyze into 50 mM Tris pH 7.5, 200 mM NaCl, 12.5 mM EDTA in order to remove any residual Ni^2^+.
6. To regenerate a column for subsequent use, strip the resin with 4-5 CVs of 100 mM EDTA; remove the EDTA by washing with 5-6 CVs of dH_2_O (and, for long-term storage, wash with 20% EtOH).
12. To proteolytically remove the His6×-tag (*see* **Note 5)**, incubate the sample overnight with thrombin at a 1:600 mass ratio of thrombin:Hfq.
13. To remove thrombin from the latest sample, either apply the material to a benzamidine column or mix it with a resin (in batch mode).
14. Additional chromatographic steps, such as size exclusion chromatography (Fig 2), may be necessary in order to isolate the various populations of hexameric, ‘free’ Hfq versus any subpopulations with RNA bound.

### 3.2 Large-scale Synthesis and Purification of the sRNA

In addition to purified protein, crystallizing an RNP complex also requires milligram quantities of RNA of sufficient quality. Here, ‘quality’ means that the ideal RNA sample will be (i) *chemically uniform*, in terms of sequence, length, and phosphate end-chemistry (i.e., uniform covalent structure), and also (ii) *structurally homogeneous* (i.e., narrow distribution of conformational states in solution). The first issue—monodispersity—is a fairly straightforward matter of chemistry, and is within one’s control (e.g., use an RNA synthesis scheme that minimizes heterogeneity of the 3′-termini of the product RNA molecules). The second issue, concerning structural heterogeneity, is a matter of physics: one can anneal RNAs by heating/cooling, adjusting pH, ionic strength, etc., to try and modulate the solution-state behavior of an RNA, but the intrinsic structural/dynamical properties of RNA are generally not easily regulated; ultimately, one must empirically monitor the RNA and its properties of interest (e.g., ‘crystallizability’).

For sRNAs, which range from ≈50 to 200 nt, a sufficient quantity of material can be readily synthesized via run-off transcription, *in vitro*, using phage T7 RNA polymerase and a linearized plasmid as the DNA template (*see* **Note 6**). Traditional *in vitro* transcription is limited by the fact that T7 polymerase strongly prefers guanosine at the 5′-end of the transcript [56], thus adversely affecting yields for target RNAs lacking a 5′ G. In addition, the polymerase typically incorporates a few nucleotides at the 3′-end of the transcript in a random, template-independent manner, giving an RNA population that is heterogeneous in length and 3′ sequence. Both of these limitations can be avoided by including, 5′ and 3′ to the RNA sequence of interest, a pair of *cis*-acting, self-cleaving ribozymes [57,58]. The flanking ribozymes ensure that the population of RNA products is accurate and chemically uniform, given the single-nt precision with which ribozymes self-cleave at the scissile bond. In principle, any self-cleaving ribozyme can be used (hammerhead, hairpin, hepatitis δ virus (HDV), etc.). In practice, an engineered 5′ hammerhead and 3′ HDV ribozyme have been found to work well [48], and impose virtually no sequence constraints on the target RNA product; some obligatory base-pairing interactions between the 5′ hammerhead ribozyme and the target RNA sequence does mean that this region will need to be re-designed for each new target RNA construct that one seeks to produce.

A DNA template suitable for the *in vitro* transcription reaction can be generated by cloning the construct into a high copy number plasmid containing the T7 promoter upstream of a multiple cloning site (MCS). The plasmid will need to be linearized using a restriction enzyme with selectivity to a site that is 3′ of the sequence of interest. The individual components for the *in vitro* transcription reaction may be prepared by the user or purchased from a manufacturer; whole kits are also commercially available (e.g., MEGAscript T7 transcription kit, Invitrogen). Large quantities of T7 RNA polymerase can be produced in-house in a cost-effective manner by using a His-tagged construct and affinity purification (similar to that described above for Hfq). In general, the concentrations of rNTPs, MgCl_2_, and T7 polymerase in the transcription reaction will require optimization for each new RNA construct/system. General guidelines and examples can be found in [57-59]. Using the method outlined below, it is ideally possible to generate milligram quantities of RNA.

1. Clone the DNA construct into a high copy number plasmid containing a T7 promoter upstream of a MCS (e.g., the pBluescript or pGEM series; *also see* **Note 7**)
2. Linearize the plasmid using a restriction enzyme for a site 3′ to the sequence of interest.
3. Mix the following components (final concentrations are noted) in the listed order, and incubate at 37°C for 1-2 h:

1. 1x transcription buffer (50 mM Tris-HCl pH 8.0, 10 mM NaCl, 6 mM MgCl_2_, 2 mM spermidine)
2. 2 mM rNTP mix
3. 10 mM DTT
4. Template DNA (≈0.05 μg/μl for a 3-kb linearized plasmid)
5. DEPC-treated H_2_O to bring to volume
6. 0.5 U/μl T7 RNA polymerase
4. To digest the original template, add RNase-free DNase I (2 U DNase I per 1 μg DNA template) and incubate for 30 min.
5. Purify the RNA by first separating on a denaturing (≈8 M urea) 10% w/v polyacrylamide gel and then excising the band corresponding to the transcript (*see* **Note 8**).
6. Add the gel slice to a tube containing 400 μl Elution Buffer and incubate for several hours at 4 °C. Centrifuge at 10000*g* for 10 min at 4°C and transfer the supernatant to a new tube.
7. Extract the RNA by phase separation with 1-2 V phenol:chloroform. Centrifuge at 10000*g* for 20 min at 4 °C and transfer the aqueous phase to a new tube.
8. Precipitate with 2-3 V ethanol.
9. Resuspend in dH_2_O or an appropriate buffer (e.g., Tris-HCl pH 7.5, 0.1 mM EDTA).

### 3.3 Crystallization of the Hfq•sRNA complex

There is no reliable way to predict the conditions that will yield well-diffracting crystals of a macromolecule or macromolecular complex, such as an Hfq•sRNA assembly: the process is almost entirely empirical. The word ‘almost’ appears in the last sentence because the process of crystallizing biomolecules is one of guided luck. Many of the biochemical properties and behavior of a system that are most salient to crystallization—idiosyncratic variations in solubility with pH, metal ions, presence of ligands, etc.— become manifest as the knowledge that one develops after many hours of working with a biomolecular sample at the bench. This implicit knowledge is highly system-specific (sometimes varying for even a single-residue mutant in a given system), it accumulates in a tortuously incremental manner, and it directly factors into the decision-making steps that ultimately dictate the success of a crystallization effort. Thus, the best advice for crystallizing an Hfq•sRNA complex is to work as extensively as possible to characterize the Hfq and sRNA components, as well as the assembled RNP, prior to extensive crystallization trials.

Ideally, one’s samples will be structurally homogeneous, thus increasing the likelihood of successful crystallization. Even given that, still it is often necessary to empirically screen through myriad potential crystallization conditions. High-throughput kits are available for the rapid screening of the myriad conditions that have successfully yielded crystals for various proteins in the past (a technique commonly referred to as sparse-matrix screening [60]). We recommend the Natrix Screen (Hampton Research), as it is specifically tailored to nucleic acid and protein-nucleic acid complexes; other commonly used screens, such as the Crystal Screen and PEG-Ion Screen are also advised. These kits are available in 15-ml or 1-ml highthroughput (HT) formats. Once a potential crystallization condition is identified, further optimization is usually required in order to improve the quality of the crystalline specimen. Often, this is pursued via ‘grid screens’. In grid screens, one or two free parameters are varied in a systematic manner; these parameters often include the buffer and pH, protein concentration, salts (types, concentrations), types and concentrations of other precipitants (e.g., PEGs), inclusion of small-molecule additives, temperature, etc. Further information on crystallization can be found in many excellent texts (e.g., [61]) and other resources, such as the Crystal Growth 101 literature available online (https://hamptonresearch.com/growth_101_lit.aspx).

In general, the purified Hfq and sRNA must be prepared and then assessed for homogeneity and stability before crystallization trials begin (often this is done via biophysical approaches or, ideally, via functional assays). Also, we advise adhering as closely as possible to RNase-free procedures (e.g., use DEPC-treated water, RNase Zap) both in biochemical characterization steps and in handling Hfq, sRNA and Hfq•sRNA specimens for crystallization trials. The following is a general protocol to get started:

1. Dialyze the purified Hfq into a suitable crystallization buffer. This should be the simplest, most minimalistic buffer in which the biomolecule is stable and soluble, to a concentration of at least 1-2 mg/ml; for instance, a buffer such as 20 mM TrisCl pH 7.5, 200 mM NaCl has often worked well in our experience. Because RNA is involved, inclusion of salts of divalent cations, such as MgCl2, may be found to aid in crystallization and overall diffraction quality.
2. Bring the [Hfq] to ≈15 mg/ml via concentration or dilution, as necessary (*see* **Note 9**); concentration is often achieved via centrifugal filtration devices with a suitable MWCO.
3. To potentially enhance the conformational homogeneity of the sRNA via annealing, incubate at 80 °C and slowly cool to RT; another approach worth trying is to heat the RNA sample and then snap-cool to ≈4°C on ice.
4. As a cautionary (and troubleshooting) step, one can test for background RNase activity in the crystallization sample by incubating the Hfq and RNA together for ≈2 weeks and assay degradation via PAGE or other methods (a molar ratio of between 1:1 and 2:1 protein:RNA is recommended as a starting point [25,48,62]).

Finally, to begin crystallization trials one should follow the manufacturer’s protocol for the particular sparse-matrix screen. Crystal trays should be stored in a temperature and humidity-controlled environment. Crystals can take between hours and months to develop. We recommend checking trays for crystals relatively frequently at the start of the process—e.g., after 1, 2, 4, 8, 16, 32 days. Experience suggests that, ideally, precipitation should occur in roughly one-third to one-half of the conditions within minutes of setting-up the crystallization drop; if this is not the case, the Hfq, sRNA, or Hfq•sRNA concentrations may need to be adjusted accordingly. Once a potential crystallization condition has been identified, it should be re-made in-house (using one’s own reagents) in order to ensure reproducibility. Then, largescale grid screening and further optimization can be pursued.

Intriguingly, a survey of the PDB identifies several crystallization agents that seem to recur in the crystallization of Hfq and Hfq•RNA complexes, as detailed in Table 1. The most commonly occurring reagents are (i) sodium cacodylate and citrate buffers, (ii) PEG 3350 and 2-methyl-2,4-pentanediol (MPD) precipitants, and (iii) MgCl_2_, CoCl_2_, and KCl salts as additives. Other divalent cations and polyamines, such as metal hexammines (e.g., hexammine cobalt(III) chloride, [Co(NH_3_)_6_]Cl_3_), spermine, and spermidine, have been found to aid in the crystallization of many protein•nucleic acid complexes [48,62].

Several methods can be applied to verify that new crystals are indeed of an Hfq•sRNA complex. An Intelli-Plate (Hampton Research, HR3-118) may be used during sparse-matrix screening in order to test, in parallel, multiple components for each crystallization condition (e.g., the Hfq•sRNA complex, the Hfq alone, and the buffer alone). If crystals appear only in the drop containing Hfq•sRNA complex, there is a high likelihood that the crystals are composed of the complex. Also, macromolecular crystals can be washed, dissolved, and run on an SDS-PAGE or native polyacrylamide gel, or one can subject them to the flame of a Bunsen burner (biomolecular crystals melt, whereas salt crystals survive this trial by fire [63]). Small-molecule dyes, such as crystal violet or methylene blue, are taken up by macromolecular crystals but not by salt crystals, and thus can be used to distinguish between the two[64]. Finally, obtaining a diffraction dataset is the ultimate way to determine if a given crystal is macromolecular and, if so, the likelihood of a successful structure determination from that specimen.

## 4 Notes

1. Hfq is known to protect RNAs [65], and nucleic acid may still remain even after nuclease treatment. This potential pitfall should be monitored by A_260_/A_280_. If protein degradation is detected by SDSPAGE or other means, then the Lysis Buffer used to resuspend the frozen cell pellet should be supplemented with a protease inhibitor cocktail, either commercial or home-made (including such compounds as PMSF, AEBSF, EDTA, aprotinin, leupeptin, etc.).
2. Experience with many recombinant Sm-like protein constructs suggests that the efficacy of the heat-cut step (i.e., degree of purification achieved) can vary greatly with temperature: we have found that many (>5) more *E. coli* proteins retain solubility in the clarified lysate after a 70 °C heat-cut, versus at 75 °C, at least for the BL21(DE3) strain. In purifying a new Hfq homolog, one can test 500 μLaliquots of the clarified lysate at a series of temperatures near this range, say 65, 70, 75 and 80 °C.
3. In our experience, Hfq withstands treatment with conventional chaotropic agents, such as high concentrations (≈6-8 M) of urea or GndCl. While such treatment may not fully denature the protein, we have found that it can disrupt potential Hfq…nucleic acid interactions. Adding such denaturants to the wash and elution buffers used in the IMAC stage can help mitigate nucleic acid contamination.
4. The divalent cation Co^2^+ has a lower affinity (than Ni^2^+) for the imidazole side-chain of histidine, but it also features less non-specific binding to arbitrary proteins; if necessary because of persistent contaminants in the Hfq eluate, one can try Co^2^+ in place of Ni^2^+ in the critical IMAC purification step.
5. Often, proteins are crystallized with an intact His6x-tag. Protein tags can potentially interfere with structure or function, although this is less likely with the small His6x-tag. His-tags can also deleteriously affect crystallizability, by increasing the length of a disordered tail or by forming spurious (and weak) crystal contacts that lead to lattice disorder. We recommend cleaving the tag if possible, as this better replicates the wild type sequence. If crystals cannot be obtained with the un-tagged protein, the tagged construct should be considered for crystallization too. As two practical anecdotes from our work with the Sm-like archaeal protein (SmAP) homologs of Hfq, we note the following:(i) with *Pyrobaculum aerophilum* SmAP1, a C-terminal His6x tag was found to interfere with oligomerization *in vitro*, and ultimately the tag was cleaved-off in order for crystallization to succeed [66], and (ii) for that same recombinant construct, attempts to remove the His-tag via treatment with thrombin failed (even though the linker between the tag and the native protein sequence was designed to include a thrombin recognition site), but the tag could be successfully removed by proteolytic treatment with trypsin. In such work, we generally use mass spectrometry (typically MALDITOF, sometimes electrospray) to assess the accuracy of the cut-site and completeness of proteolysis.
6. Previous work has examined the RNA-binding properties of Hfq using either (i) free ribonucleotides of various forms (e.g., rNTPs, rNMPs, etc.), (ii) short oligoribonucleotides of ≲30-nt, e.g. the rU_6_ oligo co-crystallized by Stanek et al. [23], or (iii) longer, full-length sRNAs, such as the 65-nt *Salmonella* RydC sRNA co-crystallized with *E. coli* Hfq [25]. The nucleotides in category (i) are readily purchased, and the oligonucleotides in category (ii) are readily obtained via stepwise, solid-phase chemical synthesis (such RNAs are available from various suppliers, e.g. Dharmacon). In contrast, such approaches are inefficient for the longer (≳30-nt) oligonucleotides of (iii), and these can be efficiently generated by enzymatic synthesis, using RNA polymerase *in vitro*.
7. The plasmids pUC18 and pUC19 are commonly used for *in vitro* transcription. The T7 promoter will need to be cloned into these plasmids as well.
8. If the RNA construct contains ribozymes, be careful to not overload gels in order to enable the correctly processed, self-cleaved RNA transcript to be isolated from other products that differ by only a few nucleotides.
9. Typically, proteins are crystallized at concentrations between ≈5-20 mg/ml. Nevertheless, concentrations well outside this range have been required for Hfq and other Sm proteins; for instance, *A. aeolicus* Hfq crystallized at 4 mg/ml [23], while *P. aerophilum* SmAP3 was at 85 mg/ml [67]. Hfq concentrations may well be limited by protein solubility (not just supply), and likely will need to be varied in any successful set of crystallization trials.
10. Tacsimate is “a mixture of titrated organic acid salts” that contains 1.8 M malonic acid, 0.25 M ammonium citrate tribasic, 0.12 M succinic acid, 0.3 M DL-malic acid, 0.4 M sodium acetate trihydrate, 0.5 M sodium formate, and 0.16 M ammonium tartrate dibasic (for more information, see http://hamptonresearch.com/documents/product/hr000175_what_is_tacsimate_new.pdf).
11. This structure contains molecules of both ADP and the non-hydrolyzable ATP analog AMP-PNP.
12. In this serendipitous co-crystal structure of *E. coli* Hfq and catalase HPII, an Hfq hexamer was found to bind each subunit of a HPII tetramer.
13. MPEG, an acronym for methoxypolyethylene glycol (also known as PEG monomethyl ether), has a covalent formula of CH_3_(OCH_2_CH_2_)*n* OH, versus H(OCH_2_CH_2_)*n* OH for simple PEGs.

## Acknowledgements

We thank L. Columbus (UVa) for helpful discussions. This work was funded by NSF CAREER award MCB–1350957.

## Figure Legends

**Figure 1** Crystal structure of Hfq in complex with RydC sRNA (PDB 4V2S) (A) The sequence of *S. enterica* RydC sRNA is shown. The grey residues were not discernible in the crystal structure and were manually modelled in (B) and (C). Residues that bind Hfq at the lateral and proximal sites are highlighted. (B) In this cartoon ribbon representation of the *E.coli* Hfq hexamer, alternating monomeric subunits are colored blue and cyan. N′and C′-termini are labelled for the monomer at the 6-o’clock position. The RydC RNA backbone is shown as a tan-colored tube, with the termini labelled. The 3′ end of the RydC RNA wraps around the *proximal* pore of the Hfq ring, and an internal region of the RNA binds to the *lateral* rim (yellow arrow). Uracil bases involved in binding Hfq at the *proximal* and *lateral* sites are thickened and colored orange and yellow (respectively). (C) The RydC sRNA mediates crystal contacts via binding to the *lateral* pocket of an adjacent Hfq hexamer, as indicated by the red arrow. The same coloring scheme is used as in (B), with the uridines that facilitate crystal contacts thickened and colored red. This figure was created with PyMOL.

**Figure 2** Size-exclusion chromatography of Hfq samples reveals the impact of a chaotrope such as guanidinium chloride (GndCl) on elution profiles and co-purifying nucleic acid content. In particular, high concentrations of GndCl can disrupt Hfq…RNA interactions, as shown here via preparative-scale SEC chromatograms for recombinant His-tagged *A. aeolicus* Hfq constructs that were previously purified by IMAC either in the absence (0 M) or presence (at 3 M, 6 M) of GndCl. The peak that elutes at ≈60 ml corresponds to Hfq associated with nucleic acids, as indicated by the higher molecular weight (versus Hfq alone) and the high A_260_/A_280_ absorbance ratio for this eluate; the peak at ≈100 ml corresponds to pure Hfq protein. Note the smooth shift from nucleic acid–bound Hfq to free protein as [GndCl] increases.

